# ATRX promotes transcription initiation of HSV-1 immediate early genes during early lytic infection

**DOI:** 10.1101/2025.04.14.648792

**Authors:** Laura E.M. Dunn, Mackenzie M. Clark, Joel D. Baines

## Abstract

Herpes simplex virus 1 (HSV-1) transcribes its genome in a highly coordinated temporal cascade, utilizing the host RNA polymerase II (Pol II). Repression of transcription precedes the cascade’s progression in a process requiring viral immediate early (IE) genes, in a phenomenon termed Transient Immediate Early gene Mediated Repression (TIEMR). Given that components of promyelocytic leukaemia nuclear bodies (PML-NBs) are known to rapidly engage incoming HSV-1 genomes, we investigated potential roles in TIEMR regulation. Using siRNA knockdown of PML-NB constituents (PML, DAXX, and ATRX), we observed that ATRX depletion significantly reduced nascent viral transcription on viral IE promoters at 1.5 hours post-infection (hpi) while DAXX knockdown increased viral transcription. ChIP-Seq analysis indicated ATRX associates with both highly transcriptionally active viral IE genes and with transcriptionally restricted non-IE viral genes, suggesting diverse functions. We show that ATRX’s association with active transcription on IE genes correlates with the presence of G-quadruplexes (G4s). Pharmacologic stabilization of G4s mimicked the effects of ATRX knockdown, significantly reducing transcription initiation on IE genes. These findings suggest that ATRX promotes transcriptional initiation of HSV-1 IE genes by preventing or ablating G4 formation. Our results reveal a previously unrecognized pro-transcriptional role for ATRX early in HSV-1 infection.

## Introduction

At the initiation of HSV-1 infection, histone-free viral genomes encoding more than 80 densely packed genes are delivered to the nucleus. These genomes contain a high concentration of promoter elements, including TATA boxes and initiator sequences recognized by cellular RNA Polymerase II (Pol II) (1, 2). As such, the viral genome is engaged by Pol II within minutes of infection (3, 4). However, the virus rapidly represses Pol II activity, primarily through its own immediate early (IE) proteins, in a process known as Transient Immediate Early gene Mediated Repression (TIEMR) (5). Once this repression is established, transcription of viral genes is regulated by de-repression, leading to a temporal cascade of expression of different gene subsets defined as IE, early (E), leaky late (LL), and true late (L) (6, 7).

Many nuclear proteins engage the entering HSV-1 DNA within the first hour of infection. These include proteins involved in transcription, RNA processing, chromatin regulation, and the DNA damage response (8). Multiple laboratories have also shown that nuclear HSV-1 genomes rapidly colocalize with Promyelocytic Leukaemia Nuclear Bodies (PML-NBs) (8–11). PML-NBs are multi-protein nuclear domains containing core components PML, sp100, ATRX, and DAXX (11, 12). The sequestration of viral DNA in PML-NBs is assumed to be an intrinsic immune response to repress HSV-1 infection (13–15). HSV-1 counteracts this response through the PML-NB antagonist, ICP0, which acts as a RING-finger ubiquitin ligase to promote the degradation of PML, resulting in the dissolution of PML bodies (16, 17).

The requirement for early transcriptional repression on the viral genome (5) led us to hypothesize that the virus exploits PML-NB-associated repression to facilitate the establishment of TIEMR. Here, we demonstrate that while DAXX does contribute to early transcription repression, we unexpectedly found that its binding partner (18) ATRX promotes transcription initiation on viral IE genes at 1.5 hpi. ATRX has traditionally been regarded as a repressor due to its stabilizing H3.3 on the viral genome, which can be linked to heterochromatin formation (19, 20). However, using a ChIP-Seq protocol that improves the detection of proteins such as ATRX that bind DNA indirectly, (21) we show that, as on different cellular genes, ATRX localizes not only to restricted transcription regions but also actively transcribing sites, suggesting a more complex and diverse role in ATRX regulation of viral gene expression than previously appreciated. Furthermore, we provide evidence supporting the hypothesis that ATRX-mediated transcriptional activation is facilitated by its association with G-quadruplex (G4) DNA structures at viral IE promoters.

## Results

### ATRX depletion reduces transcriptional activity on HSV-1 genes at 1.5 hpi

To dissect the roles of individual PML-NB constituents in regulating early HSV-1 transcription, we first developed a system to screen for the effects of gene knockdowns on nascent transcription. Previous work in our laboratory relied on nuclear run-on assays coupled with sequencing (3, 22). Though this method provides unparalleled detail of Pol II activity, it is time-consuming and requires a large amount of starting material (0.5-2×10^7^ cells), making the protocol unsuitable for screening. We adapted an RT-qPCR-based method (23) to address this challenge by incorporating biotin-NTP run-on and streptavidin pulldown of nascent RNA. HSV-specific primers were then used for RT-qPCR to assess transcriptional activity at specific regions on the HSV-1 genome, normalized to the human gene ACTB (validated as a run-on reference gene by Roberts et al. (23)). An overview of this protocol is shown in Fig. 1A. Primers were designed to amplify regions known to have detectable levels of nascent transcription on the viral genome at 1.5 hours post-infection (hpi) (5), (Listed in Table S1). Knockdown was achieved through siRNA transfection of HEp-2 cells, targeting the PML-NB constituent genes PML, DAXX, and ATRX alongside a non-targeting negative control. After 48h, cells were infected with HSV-1 (F) at an MOI of 5, and nuclei were harvested at 1.5 hpi. RT-qPCR confirmed at least 80% knockdown at the mRNA level of each target gene relative to the negative control (Fig. 1B). Nuclear run-on was then performed, and RT-qPCR was used to quantify nascent transcription of *UL2, UL54 (a27)*, and *US1(a22)* on the viral genome. The run-on qPCR data was additionally corrected by viral genome copy number, which was determined via qPCR on DNA isolated from the respective samples. HSV-1 genome copy results are shown in Fig. 1C, indicating only minor variations in copies of input viral genomes among siRNA treatments.

**Figure 1:**
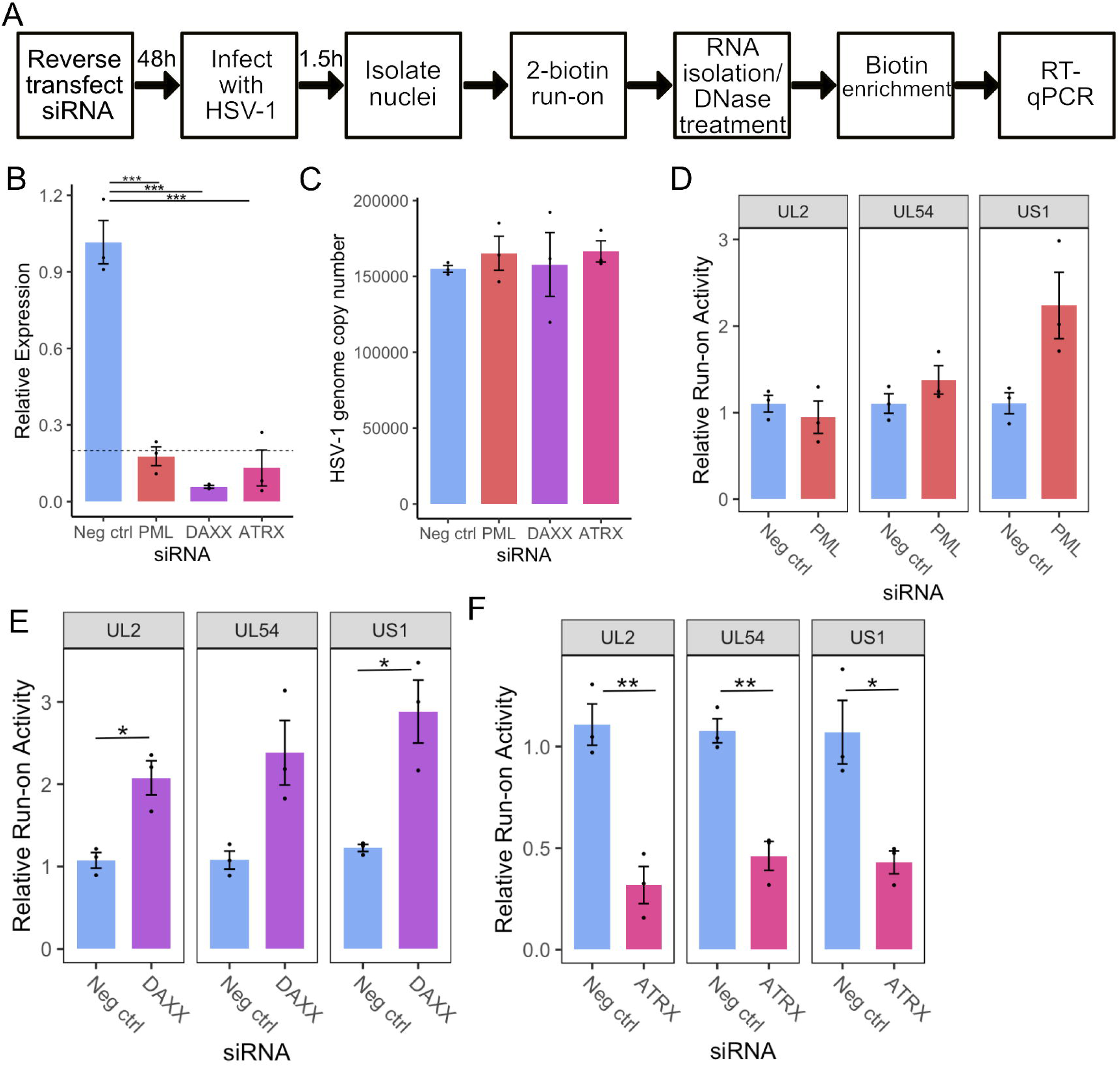
ATRX depletion reduces transcriptional activity on HSV-1 genes at 1.5 hpi. A) Overview of siRNA KD coupled to RT-qPCR methodology for nascent RNA analysis. B) RT-qPCR validation of siRNA knockdown at the mRNA level. Data are mean ± standard error. Plotted values are relative to the average negative control (non-targeting), normalized to SDHA. Statistical significance was determined using Dunnett’s test, with neg ctrl as the control group. C) HSV-1 genome copy per 5 ng of DNA in siRNA knockdown at 1.5 hpi, determined by UL51 plasmid standard curve qPCR. Data are mean ± standard error. Nascent transcriptional activity of HSV-1 genes at 1.5 hpi during PML knockdown (D), DAXX knockdown (E), and ATRX knockdown (F). Data are mean ± standard error. Plotted values are relative to the average negative control (non-targeting), normalized to ACTB and viral genome copy number. Statistical significance was determined using Welch’s t-test. Asterisks indicate statistical significance (*=p<0.05, **=p<0.01, ***=p<0.001).

Depletion of PML had a minimal effect on viral transcription, with only a detectable increase on US1 that did not reach statistical significance (Fig. 1D). Depletion of DAXX led to an approximately 2-fold increase of transcriptional activity at all regions analyzed (Fig. 1E). In contrast, depletion of DAXX’s binding partner, ATRX, resulted in an approximately 2-fold decrease in transcription of all genes tested (Fig. 1F). We confirmed these changes in nascent transcription during ATRX or DAXX depletion in TERT-immortalized human foreskin fibroblasts (HFFs). As in HEp2 cells, DAXX knockdown in HFFs increased *UL2, UL54*, and *US1* transcription at 1.5 hpi, whereas ATRX knockdown reduced transcription of these genes (Fig. S1). Thus, it is apparent that ATRX and DAXX have distinct roles in early transcriptional regulation and that ATRX promotes transcription upon initial infection.

### PML-NBs retain the ability to form in ATRX-depleted cells

The finding that ATRX plays a pro-transcriptional role was interesting because this contrasts with its assumed role as a restriction factor (24). We, therefore, sought to characterize this activity in more depth using our siRNA system. First, RT-qPCR confirmed that the loss of nascent transcription also reduced HSV-1 mRNA (Fig. S2A). As DAXX and ATRX are binding partners (18), we also assessed DAXX expression by RT-qPCR during ATRX KD, which revealed no significant effect on DAXX mRNA levels (Fig. S2B).

Next, we analyzed the ability of PML-NBs to form in the absence of ATRX to ensure the effect was not caused indirectly by the perturbation of PML-NBs. ATRX-depleted cells were stained for both PML and ATRX by immunofluorescence. PML foci were distinct in the Neg ctrl and showed strong colocalization with ATRX foci (Fig. 2A, upper panel). While the ATRX KD cells displayed distinct PML foci, they lacked detectable ATRX foci, as expected (Fig. 2A, lower panel). CellProfiler software (25) was then used to quantify these observations. ATRX KD did not significantly affect the PML foci counted per cell (Fig. 2B) but confirmed a significant depletion in ATRX foci per cell (Fig. 2C). In addition, over 80% of the PML-positive cells from ATRX KD contained no detectable ATRX foci (Fig. 2D), further indicating that the siRNA knockdown of ATRX mRNA conferred a decrease in ATRX protein in PML-NB’s.

**Figure 2:**
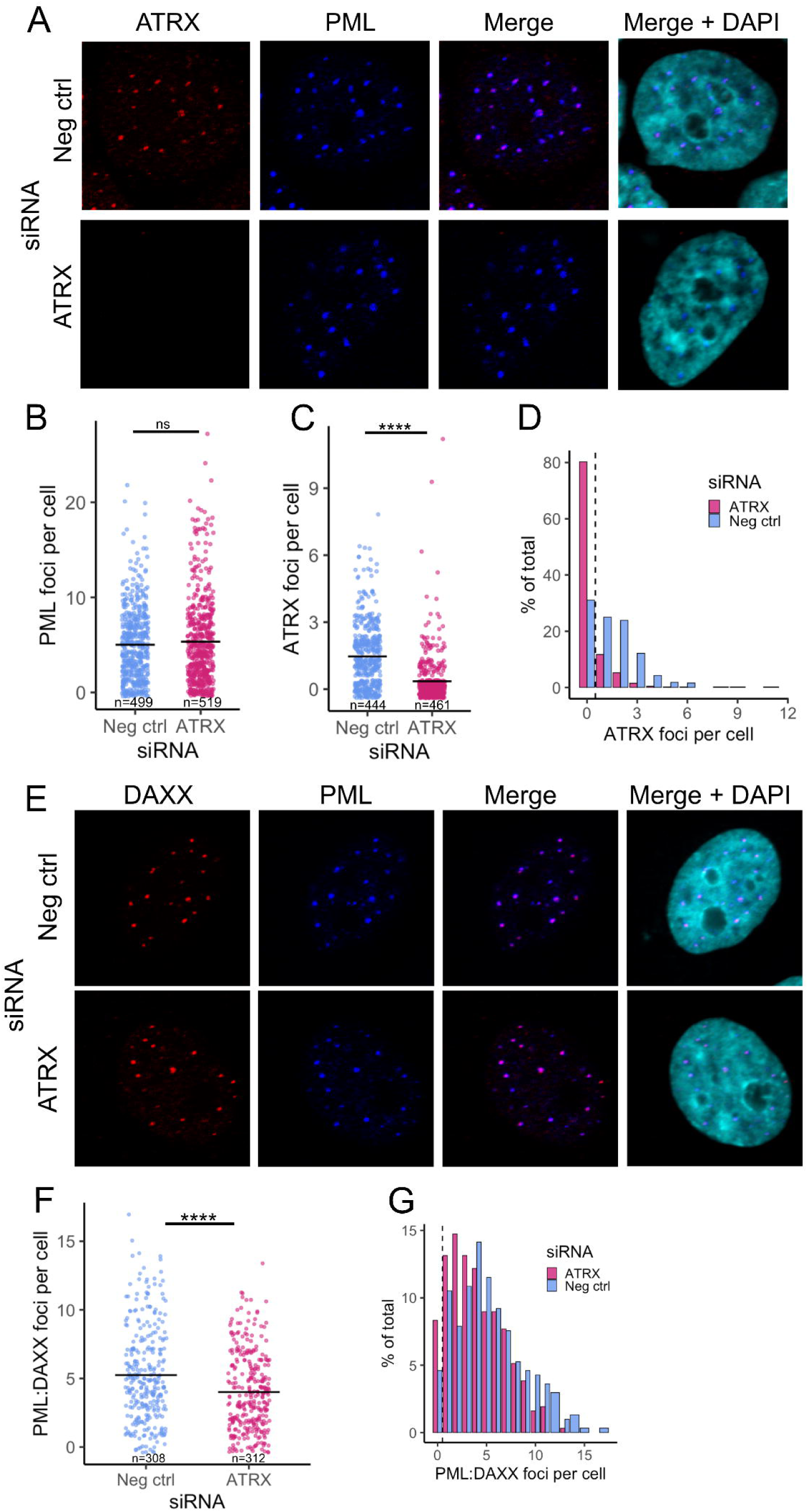
PML-NBs retain the ability to form in ATRX-depleted cells. A) Representative confocal images of ATRX (red) and PML (blue) expression after 48h ATRX siRNA knockdown, compared to neg ctrl (non-targeting) knockdown. Nuclei were stained with DAPI. Quantification of PML (B) and ATRX (C) foci per cell (nucleus). Each dot represents an individual cell. n = number of nuclei counted per condition. D) Histogram of the frequency of ATRX foci counts. E) Representative confocal images of DAXX (red) and PML (blue) expression after 48h ATRX siRNA knockdown, compared to neg ctrl (non-targeting) knockdown. Nuclei were stained with DAPI. F) Quantification of the PML:DAXX colocalized foci per cell. Each dot represents an individual cell. n = number of nuclei counted per condition. G) Histogram of the frequency of PML:DAXX foci counts. Statistical significance was determined using Welch’s t-test. Asterisks indicate statistical significance (****=p<0.0001).

The unchanged total PML foci per cell in ATRX KD indicated that PML-NBs still form. To further validate this, we also stained for DAXX, another major component of PML-NBs and a binding partner of ATRX. Both PML and DAXX foci were readily detectable. They appeared to colocalize in both ATRX KD and Neg ctrl (Fig. 2E). CellProfiler was then used to quantify the number of DAXX foci per cell (Fig. S2D) and those that colocalized with PML. This revealed a slight drop in DAXX-PML-associated NBs during ATRX KD, with the mean number dropping to 4 per cell from 5.25 in the Neg ctrl (Fig. 2F).

Plotting the distribution of DAXX/PML foci highlighted the trend toward a lower number of foci per cell during ATRX depletion. However, over 91% of cells still contained DAXX-PML-associated NBs (Fig. 2G). In addition, we saw no evidence of an increase in these repressive factors (DAXX and PML) that might contribute to the transcriptional reduction that occurs during ATRX depletion. Together, these data indicate that the transcriptional effects of ATRX knockdown are ATRX-specific and not a result of alteration of other PML-NB components.

### ATRX promotes transcription initiation on IE genes at 1.5hpi

Next, we used PRO-Seq to characterize high-resolution transcriptional activity across the viral genome in ATRX-depleted cells. HEp-2 cells were transfected with siRNA for 48h, followed by infection with HSV-1 at an MOI of 5 PFU/cell. After 1 h adsorption at 4°C, infection was allowed to proceed at 37°C. Nuclei were then harvested at 1.5 hpi, and PRO-Seq was performed. Knockdown was validated by RT-qPCR (Fig. 3A), and qPCR was used to assess input viral genome copy in the run-on nuclei, confirming there was no significant difference between groups (Fig. 3B). Noted minor differences in genome copy number were accounted for in normalization of the PRO-Seq data, which was also corrected to the total library read count to account for variation in sequencing depth (details in Table S3). Calculation of the density of reads across each HSV-1 gene indicated that ATRX depletion led to a significant reduction in transcriptional activity of all viral genes at 1.5hpi (Fig. 3C), confirming the result from the prior qPCR run-on assay (Fig. 1G).

**Figure 3:**
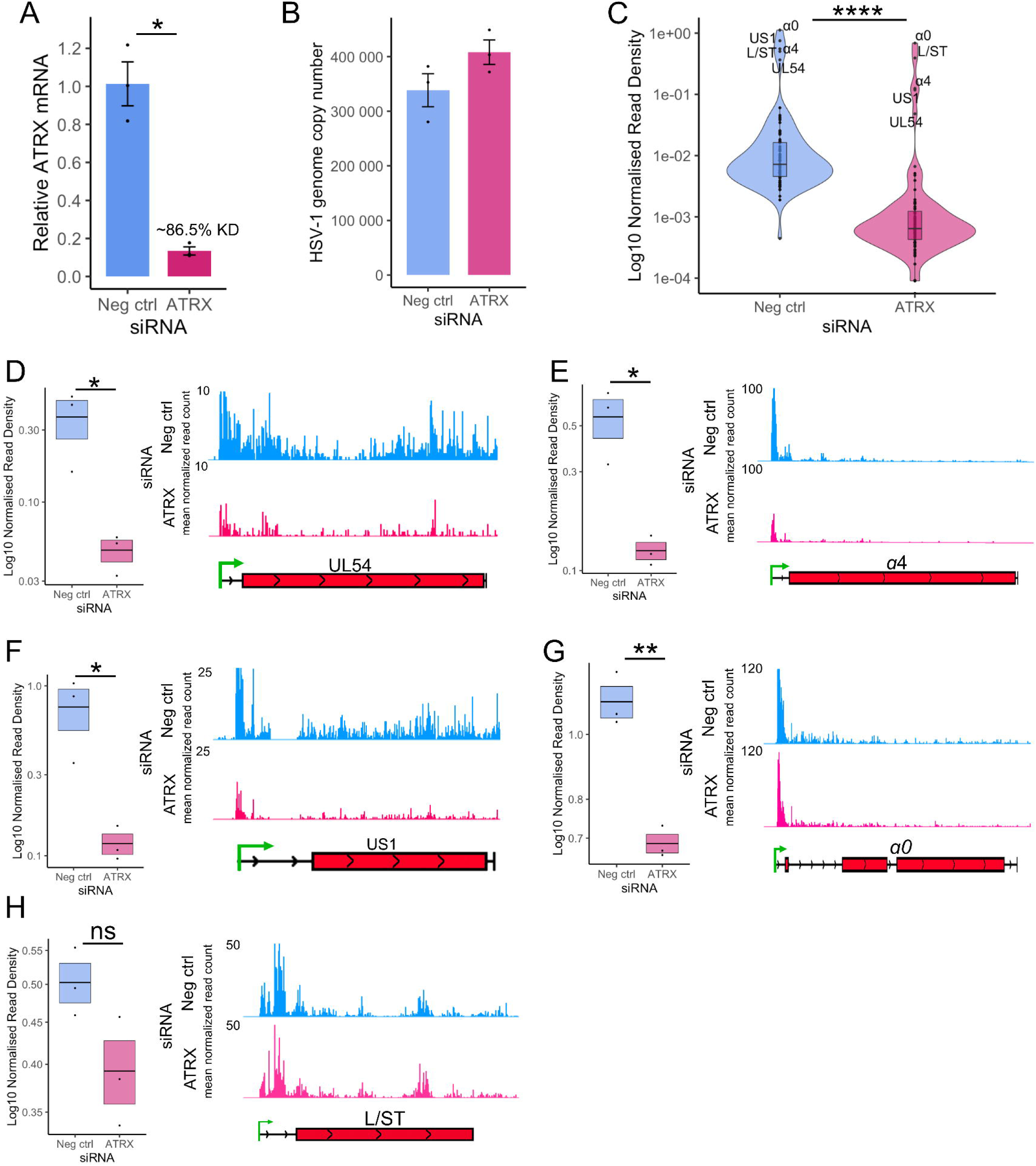
ATRX promotes transcription on IE genes at 1.5hpi. A) RT-qPCR validation of ATRX knockdown at the mRNA level in cells used for PRO-Seq. Data are mean ± standard error. Plotted values are relative to the average negative control (non-targeting), normalized to SDHA. B) HSV-1 genome copy per 5 ng of DNA in siRNA knockdown at 1.5 hpi, determined by UL51 plasmid standard curve qPCR. Data are mean ± standard error. C) PRO-Seq read density across viral genes at 1.5 hpi during ATRX knockdown, compared to neg ctrl (non-targeting) knockdown. Statistical significance was determined using the Wilcoxon test. PRO-Seq read density of individual IE genes and genome browser tracks at 1.5 hpi of D) UL54, E) α4, F) US1, G) α0 and H) L/ST. PRO-Seq read density is calculated as read per bp, normalized to spike-in and HSV-1 genome copy. Black lines indicate mean and bands ± standard error. Statistical significance was determined using Welch’s t-test. Asterisks indicate statistical significance (*=p<0.05, **=p<0.01, ***=p<0.001, ****=p<0.0001).

Next, we focused the analysis on the IE genes, the most critical genes at 1.5 hpi and the most robustly transcribed (Fig. 3C). The PRO-Seq read density of each IE gene was calculated and analyzed, alongside IGV genome browser displays for visualization (Fig. 3D-G). This confirmed a significant decrease in transcriptional activity across all IE genes. L/ST was also analyzed because it was highly transcribed at 1.5 hpi. Transcriptional activity across L/ST was not significantly altered by ATRX depletion (Fig. 3H).

Genome browser views of PRO-Seq data can obscure details due to the predominant peak near the promoter. Therefore, a metaplot of the cubic spline of PRO-Seq reads across IE genes was also plotted, highlighting the overall loss of transcriptional activity during ATRX depletion (Fig. 4A). This reduction was apparent across the whole gene but was most striking at the promoter region (inset), the site of Pol II pausing. A reduction in a promoter peak is characteristic of a decrease in initiation (26), though it can also be due to an increased release of Pol II from pausing. To assess whether the loss of activity at the promoter was related to Pol II pausing and release, the relative activity across IE genes was calculated to measure the overall distribution of Pol II activity across genes, independent of read count. This revealed no difference in the ATRX KD compared to repair (Fig. 4B), suggesting promoter-proximal pausing was unaltered. This was further confirmed by pause index analysis (ratio of read density in promoter v read density in gene body) on individual IE genes (Fig. 4C).

**Figure 4:**
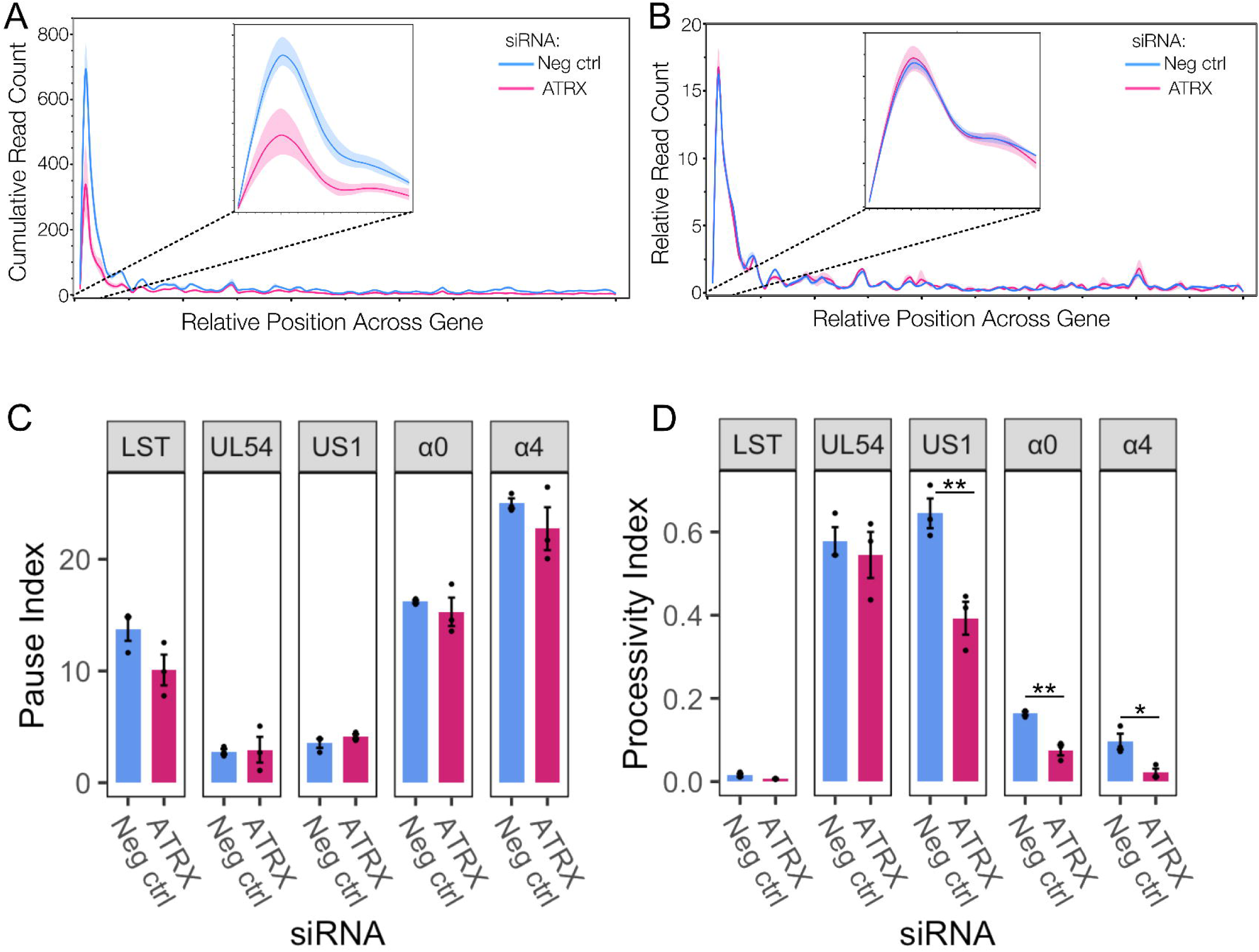
ATRX functions through promoting initiation and does not affect Pol II pausing. A) Metaplot spline interpolation of cumulative PRO-Seq read count across HSV-IE genes during ATRX knockdown at 1.5 hpi. B) Metaplot spline interpolation of relative read counts across HSV-IE genes during ATRX knockdown at 1.5 hpi. The bootstrap confidence of fit is shown in the shaded area. C) Pause and processivity index values (D) of HSV-1 IE genes during ATRX knockdown at 1.5 hpi. Data are mean ± standard error. Statistical significance was determined using Welch’s t-test. Asterisks indicate statistical significance ((*=p<0.05, **=p<0.01).

We have previously shown that Pol II is highly processive across viral genes, and extensive transcription continues beyond the polyA site in a process tightly regulated by the virus (3, 27). A processivity index was calculated to quantify Pol II processivity (ratio of read density 1000nt downstream of polyA site v read density in the gene body (27)). This revealed that specific IE genes (*US1, α0*, and *α4*) exhibit a reduction in Pol II processivity during ATRX depletion (Fig. 4D), thus indicating a decrease in the amount of Pol II that successfully reaches and proceeds beyond the polyA site. In summary, these data suggest that ATRX promotes both transcription initiation and elongation efficiency of IE genes at 1.5 hpi.

### ATRX is associated with sites of both active and repressed transcription on the cellular and viral genomes

The finding that ATRX promotes transcription on viral genes contrasts with its proposed role as a repressor. The repressive activity of ATRX is believed to result from its binding to the viral genome to promote repressive heterochromatin (19, 20). However, the high levels of transcriptional activity on the viral genome detected by this PRO-Seq study and others (3, 5, 22, 27, 28) is not characteristic of a heterochromatin state. Recent advancements in ChIP using ethylene glycol-bis(succinimidylsuccinate) (EGS) to preserve indirect and/or hyperdynamic interactions (21) have shown that ATRX is associated with both euchromatin and heterochromatin on the cellular genome (29). We, therefore, chose to use this enhanced ChIP-Seq protocol for in-depth characterization of ATRX binding on the viral genome during lytic infection. HEp-2 cells were infected with HSV-1 at an MOI of 5 PFU/cell before EGS/formalin fixation, and ChIP was then performed.

To validate the success of the ChIP-Seq, we first examined the cellular peaks (called by MACS3 (30) and quantified PRO-Seq reads (taken from an independent, non-siRNA experiment) at these peaks to assess the transcriptional activity at ATRX binding sites. As with previous EGS ARTX-ChIP studies (29), peaks were called at both intragenic (gene body and promoter) and intergenic regions, with the highest proportion found in the gene body (Fig. 5A). Notably, ATRX binding at cellular promoters correlated with significantly higher levels of transcriptional activity than intergenic regions (Fig. 5B). This is consistent with previous findings indicating that while ATRX is commonly associated with silencing at interstitial loci (31), it is also enriched at promoter regions associated with active transcription (29). An example of an ATRX peak at a promoter of a highly transcriptionally active gene is shown for ZNF131 in Fig. 5C. An example of an ATRX peak at a transcriptionally restricted region is shown in Fig. 5D; note that this region consists of simple (tandem) repeats, sequences with which ATRX is known to associate (32). In summary, the cellular gene analysis confirms the EGS ChIP methodology’s success in detecting ATRX binding at both heterochromatin and euchromatin regions, as expected. A list of sites of peak binding is given in Supplementary Data 1.

**Figure 5:**
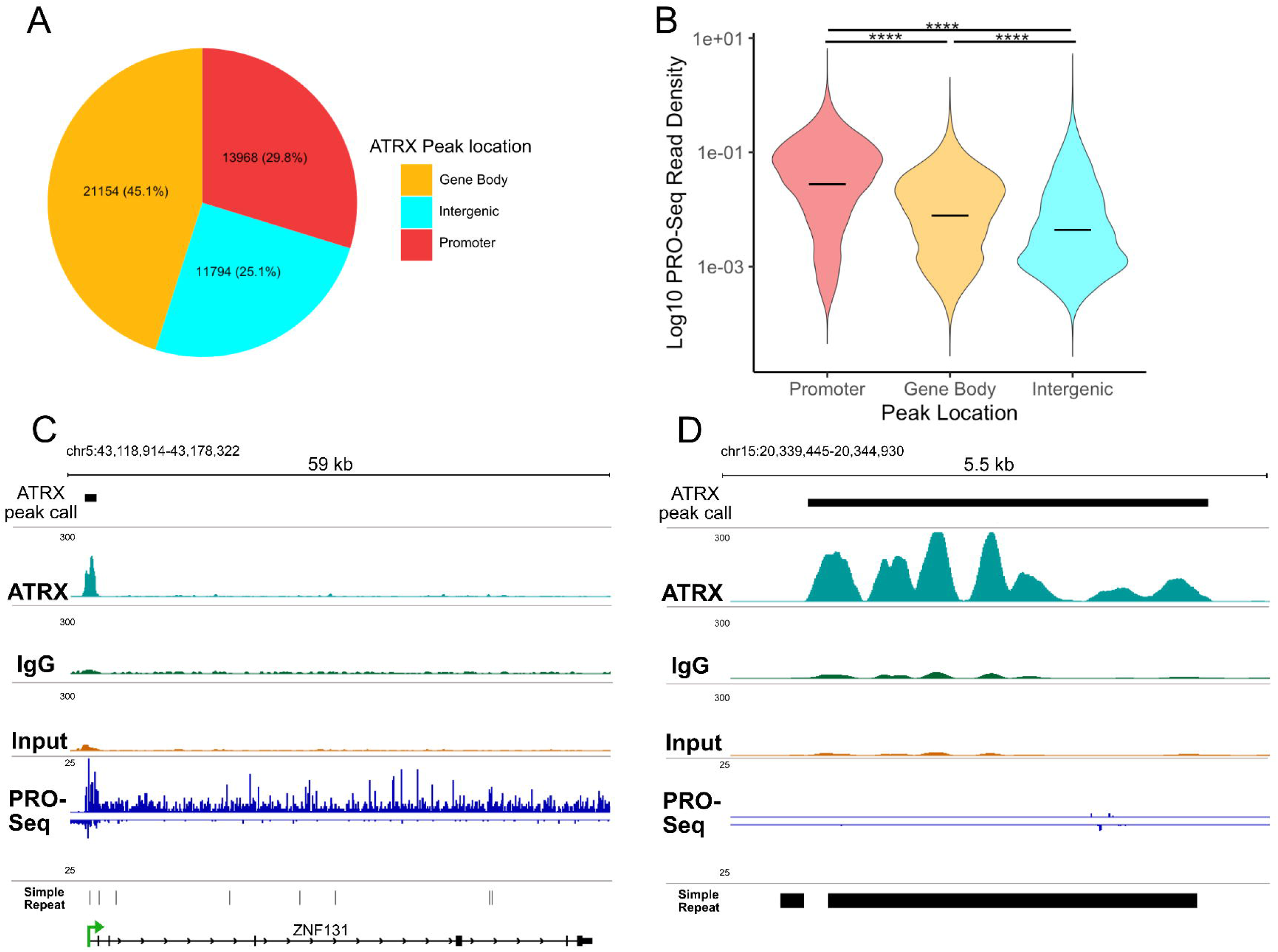
ATRX is associated with active and repressed transcription sites across the cellular genome. A) Distribution of ATRX binding sites on the cellular genome from ATRX ChIP-Seq in HSV-1 infected HEp-2 cells at 1.5 hpi. Binding sites were identified using MACS3 peak caller B) PRO-Seq read density at ATRX-binding sites (read per bp), split by peak location. The black line represents the mean. Statistical significance was determined using Wilcoxon multiple pairwise comparisons with Bonferroni correction. Asterisks indicate statistical significance (****=p<0.0001). C) Genome browser example of an ATRX peak at the promoter of the transcriptionally active cellular gene, ZNF131. D) Genome browser example of an ATRX peak across a transcriptionally silent intergenic region, containing tandem repeats. Data is the mean of 3 replicates. Genome browser coverage files were normalized to human RPGC (reads per genomic content).

The alignment of the ATRX ChIP to the HSV-1 genome is shown in Fig. 6A. As the viral genome is primarily free of tightly bound histones during lytic infection (8, 33, 34), there is the potential for unspecific IgG pulldown of viral DNA. Therefore, a stringent two-step MACS3 analysis was performed in which peaks were initially called for ATRX v Input and IgG v Input. Differential peak analysis was then applied, using a likelihood ratio to identify ATRX peaks enriched relative to IgG. The dREG tool was used on the PRO-Seq data to determine transcriptional regulatory elements (TREs) associated with accessible chromatin (35). This revealed that ATRX was enriched across the viral genome at 1.5 hpi at both highly transcriptionally active and accessible IE genes as well as transcriptionally restricted sections of the genome (Fig. 6A). To quantify these observations, ATRX binding peaks were categorized according to whether they overlapped IE or non-IE genes. This confirmed no difference in ATRX enrichment between IE and non-IE genes (Fig. 6B); however, the PRO-Seq read density in ATRX peaks was significantly higher for those that overlapped IE genes (Fig. 6C). Overall, this analysis indicated ATRX binding on the viral genome during early lytic infection is associated with both highly active and restricted transcription.

**Figure 6:**
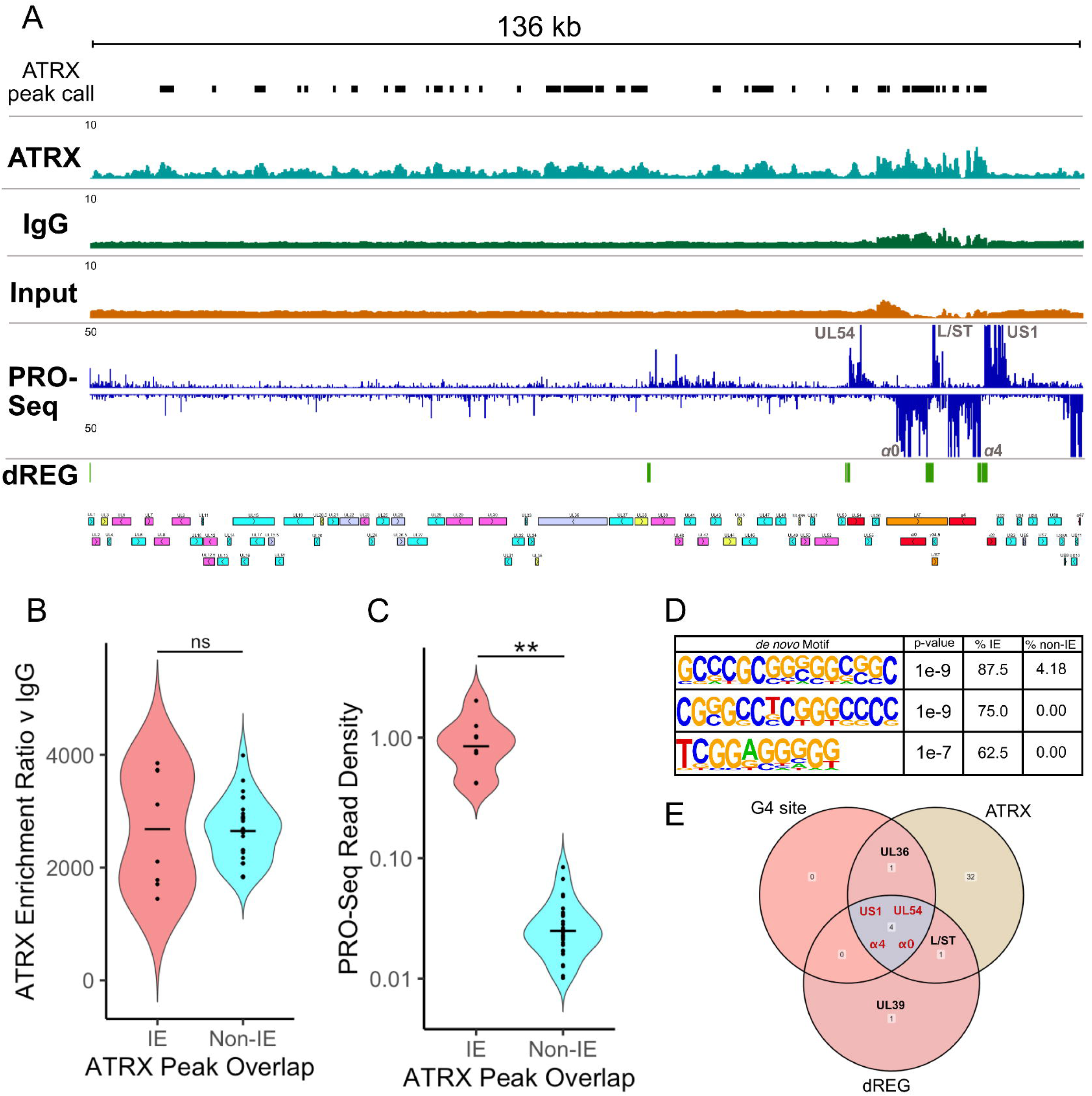
ATRX is associated with active and repressed transcription sites on the viral genome. A) Genome browser view of ATRX ChIP-Seq data across the HSV-1 F genome at 1.5 hpi with MACS3 peak calls detailed (mean of 3 replicates). HSV-1 ChIP-Seq coverage files were normalized to HSV-1 RPGC (reads per genomic content). PRO-Seq data (mean of 2 replicates) is also shown, with dREG identification of transcriptionally regulatory regions. B) MACS3 likelihood enrichment ratio of ATRX peaks, split by whether they overlap IE ornon-IE genes. C) PRO-Seq read density at ATRX peaks, split by whether they overlap IE or non-IE genes. The black line represents the mean. Statistical significance was determined using the Wilcoxon test. Asterisks indicate statistical significance (**=p<0.01). D) Enriched motifs in ATRX peaks that overlap IE genes, relative to non-IE genes. E) Overlap between viral genes, which contain G-quadruplex (G4) sites, ATRX peak, and active transcription (dREG).

This differential activity in transcription at ATRX binding sites suggests there are genomic features that might impact ATRX function. To investigate this possibility, HOMER (36) motif enrichment was used to identify sequence motifs enriched in ATRX peaks at transcriptionally active (IE) v transcriptionally restricted (non-IE) regions. Interestingly, the top three enriched motifs contained repetitive G-rich tracts (Fig. 6D), characteristic of G4 forming sequences (37). This was intriguing because ATRX was previously shown to affect gene expression through interaction with G4 regions on the cellular genome (29, 32). In addition, the HSV-1 genome contains multiple validated G4 sites, including in the promoters of IE genes (38, 39). The link between active transcription (dREG), G4s, and ATRX binding was highlighted when viral genes that contain these features were overlapped, revealing that IE genes (*UL54*, *α0, α4*, and *US1*) contain all three features (Fig. 6E).

### Stabilization of G-quadruplexes mimics the effects of ATRX depletion on viral transcription

The finding that ATRX binding on the highly transcriptional active IE genes was also linked to the presence of G4s raised the hypothesis that ATRX depletion alters G4 formation on the viral genome, thereby restricting transcription. To investigate this, we utilized the G4-ligand, BRACO-19 (40), to stabilize G4 formation during viral infection. First, we tested nascent transcription using the PRO-qPCR method. HFF and HEp-2 cells were infected with HSV-1 at an MOI of 5 PFU/cell. After a 1 h absorption period at 4°C, infection was allowed to proceed at 37°C in media containing either 25μM BRACO-19 or DMSO. Nuclei were harvested at 1.5 hpi, and PRO-RTqPCR performed. This indicated a loss of nascent viral transcription during BRACO-19 treatment in both HEp-2 (Fig. 7) and HFF cells (Fig. S3).

**Figure 7:**
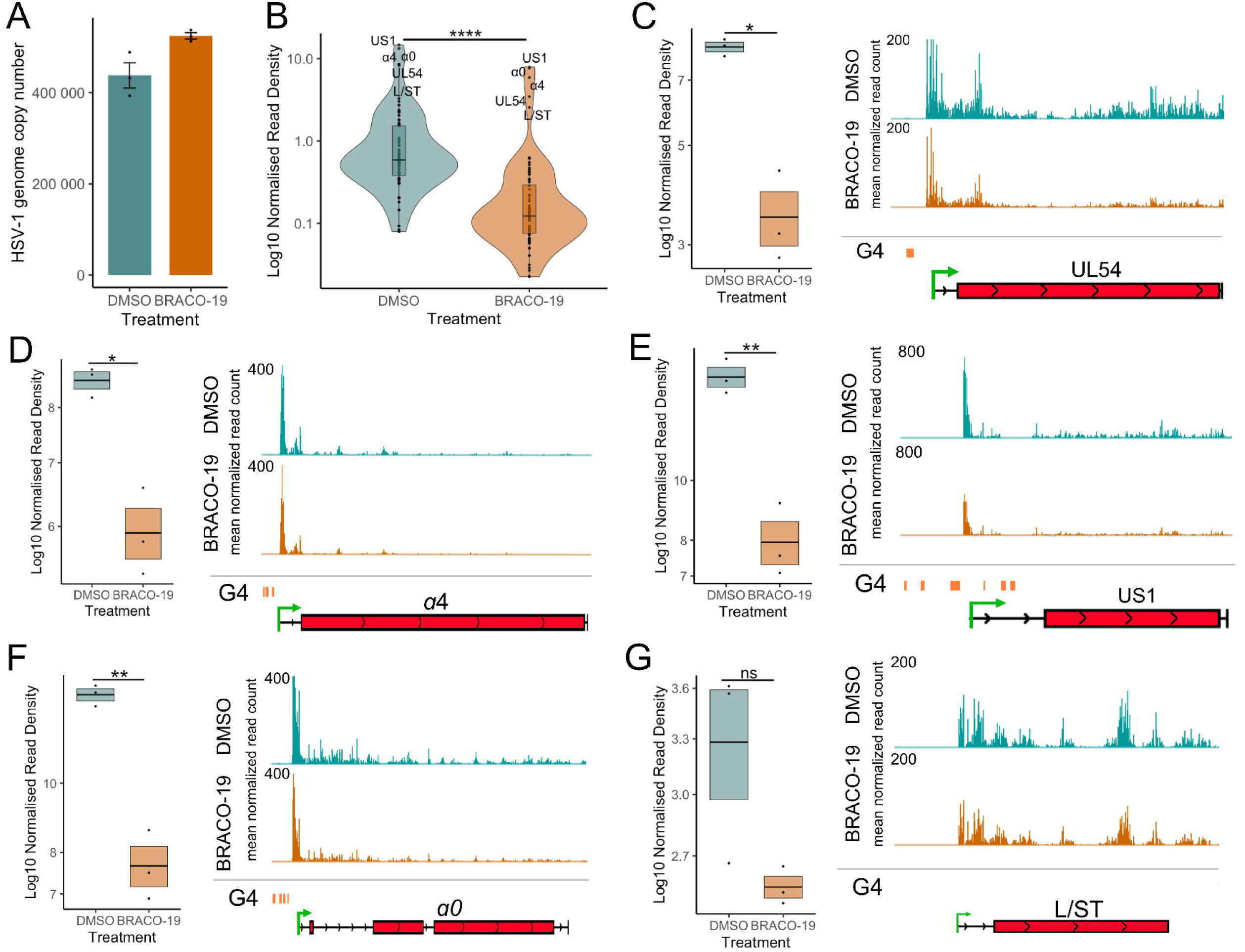
Stabilization of G-quadruplexes reduces transcriptional activity across HSV-1 genes. A) HSV-1 genome copy per 5 ng of DNA in BRACO-19 or DMSO treatment at 1.5 hpi, determined by UL51 plasmid standard curve qPCR. Data are mean ± standard error. B) PRO-Seq read density across viral genes at 1.5 hpi during BRACO-19 treatment, compared to DMSO treatment. Statistical significance was determined using the Wilcoxon test. PRO-Seq read density of individual IE genes and genome browser tracks at 1.5 hpi of C) UL54, D) α4, E) US1, F) α0 and G) L/ST. PRO-Seq read density is calculated as read per bp, normalized to spike-in and HSV-1 genome copy. Black lines indicate mean and bands ± standard error. Statistical significance was determined using Welch’s t-test. Asterisks indicate statistical significance (*=p<0.05, **=p<0.01, ***=p<0.001, ****=p<0.0001).

Next, we used PRO-Seq to analyze nascent transcription during BRACO-19 treatment in HEp-2 cells. Infections and drug treatments were performed as described above; nuclei were harvested at 1.5 hpi, and PRO-Seq was performed. qPCR was used to confirm the treatment did not affect infectivity, revealing only a slight variation in viral genome copy number between treatments (Fig. 7A). These data were included in the PRO-Seq normalization (details in Table S3). Treatment of cells with BRACO-19 led to a significant reduction in transcriptional activity across all viral genes at 1.5 hpi (Fig. 7B), similar to ATRX KD (Fig. 3C). Focusing our analysis on IE genes confirmed a significant reduction in PRO-Seq read density across all IE genes, which was also evident through IGV genome browser visualization of PRO-Seq data (Fig. 7D-F). As with ATRX KD, transcription of L/ST (which does not contain a G4 in its promoter) was not significantly affected by BRACO-19 treatment (Fig. 7G).

A metaplot of the cubic spline of the reads across IE genes was generated to quantify and compare the effect on promoter activity. This confirmed a significant reduction of transcriptional activity at IE gene promoters after BRACO-19 treatment (Fig. 8A), suggesting decreased initiation. To assess the potential for this promoter-proximal loss of activity to be a result of pause release, the relative distribution of reads was also plotted (Fig. 8B). This revealed no difference in relative transcriptional activity between BRACO-19 and DMSO treatment, indicating the effect at the promoter was not due to accelerated pause release. This was confirmed by the calculation of pause indices for all IE genes, all of which were unaffected by BRACO-19 treatment (Fig. 8C). Processivity index calculation revealed a significant reduction in Pol II processivity across all IE genes after BRACO-19 treatment (Fig. 8D), suggesting G4 stabilization restricts both initiation and efficient elongation, again similar to ATRX depletion (Fig. 4D).

**Figure 8:**
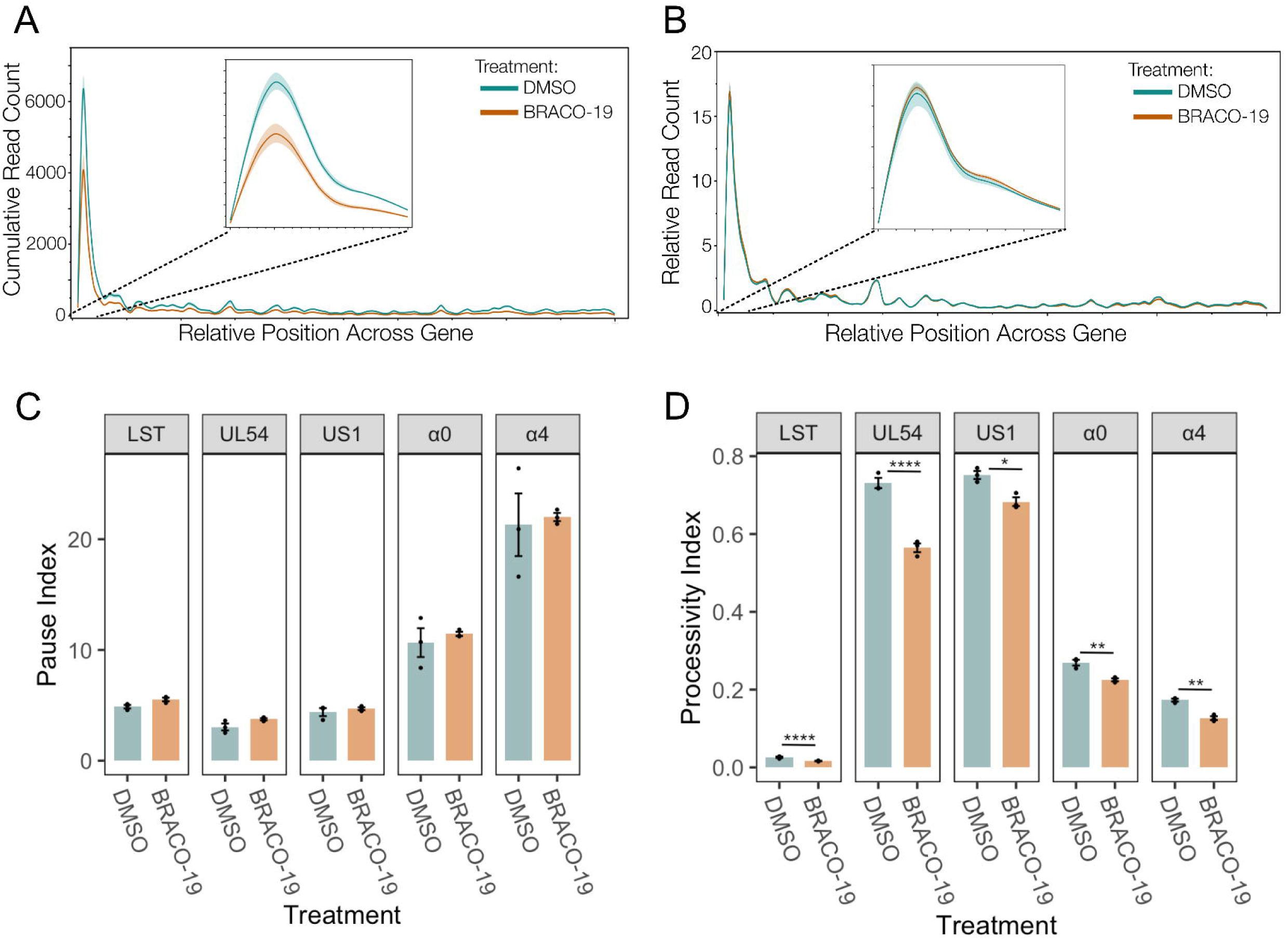
Stabilization of G-quadruplexes reduces transcriptional initiation and does not affect Pol II pausing. A) Metaplot spline interpolation of cumulative PRO-Seq read count across HSV-IE genes during BRACO-19 treatment at 1.5 hpi. B) Metaplot spline interpolation of relative read counts across HSV-IE genes during BRACO-19 treatment at 1.5 hpi. The bootstrap confidence of fit is shown in the shaded area. C) Pause and processivity index values (D) of HSV-1 IE genes during BRACO-19 treatment at 1.5 hpi. Data are mean ± standard error. Statistical significance was determined using Welch’s t-test. Asterisks indicate statistical significance ((*=p<0.05, **=p<0.01, ****=p<0.0001).

Overall, it was apparent that BRACO-19 stabilization of G4 quadruplexes mimicked the effects of ATRX KD on IE gene transcription, with decreased initiation as the predominant effect. A summary of this is shown in Figure. 9A, in which the density of PRO-Seq reads was plotted centered on IE gene promoter G4 regions. The profile of PRO-Seq reads after depletion of ATRX or BRACO-19 stabilization were almost identical at these regions, with a significant reduction in transcription at the TSS downstream of the G4 region. In summary, our data supports a model that ATRX promotes transcription initiation of IE genes through G4 destabilization. Without ATRX, G4 structures form and sterically hinder and block the initiation of transcription. A diagram of this model is shown in Fig. 9B.

**Figure 9:**
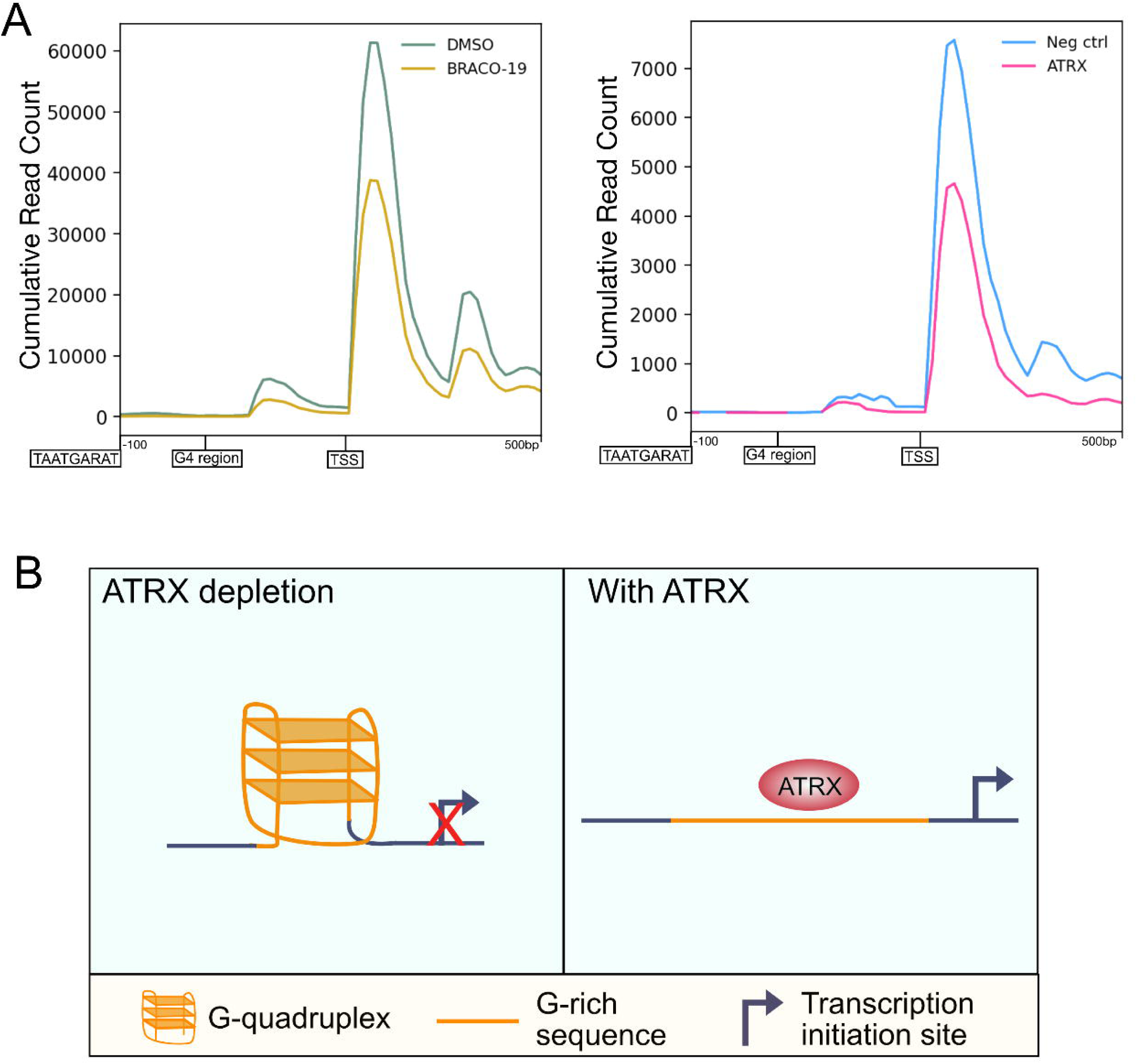
Stabilization of G-quadruplexes mimics the effects of ATRX depletion on viral transcription. Profile of PRO-Seq reads at -100 - +500 G4 regions in the promoters of HSV-1 IE genes at 1.5hpi during A) BRACO-19 treatment compared to DMSO treatment and B) ATRX knockdown compared to compared to neg ctrl (non-targeting) knockdown. B) Proposed model of the ATRX mechanism to promote transcription of viral IE genes. In the absence of ATRX, G-rich sequences in the promoters of IE genes form G4 DNA structures, sterically hindering transcription initiation. ATRX promotes destabilizing these structures to improve the accessibility of factors required for initiation.

## Discussion

Using siRNA knockdowns of PML-NB constituents and transcriptional run-on assays, we demonstrate that ATRX and DAXX, two key components of PML-NBs, play distinct roles during early viral transcription. While DAXX contributes to transcriptional repression, its binding partner ATRX unexpectedly promotes immediate early viral transcription.

The pro-transcriptional function of ATRX on the HSV-1 genome documented herein was unexpected, given prior evidence supporting a repressive function (19, 24, 41, 42). A caveat with any gene knockdown study is the potential for off-target or compensatory cellular effects, which might explain inconsistencies between experimental systems. However, we confirmed that other PML-NB components were not significantly altered (Fig. 2) and that the level of input genomes under different knockdowns remained consistent. DAXX knockdown led to increased viral transcription, in line with its known repressive role during HSV-1 infection (10, 15, 24), supporting the validity of our siRNA system.

While ATRX depletion reduced transcriptional activity across all viral gene classes (Fig. 3C), the impact on non-IE genes could be due to reduced IE protein expression. We, therefore, focused on the most highly transcribed genes at 1.5 hpi, which included IE genes and *L/ST*. Interestingly, *L/ST* transcription was unaffected by ATRX depletion (Fig. 3H). As the *L/ST* promoter lacks a VP16 transactivation motif (TAATGARAT) and G4 elements, we propose that ATRX’s function is linked to the distinct promoter architecture of IE genes.

Using PRO-Seq, which provides nucleotide resolution of transcriptional activity, we found that ATRX promotes transcriptional initiation but not Pol II pausing (Fig.4A/B). We also noted decreased Pol II processivity across specific IE genes in ATRX-depleted cells. Since Pol II activity extends beyond polyA sites before termination (43), a reduction in reads downstream of polyA signals suggests decreased elongation efficiency. This result is consistent with previous findings showing that reduced transcriptional initiation correlates with diminished Pol II elongation on HSV-1 genes (27), potentially due to decreased accessibility and impaired recruitment of elongation factors (44, 45).

ATRX has been proposed to repress transcription by stabilizing H3.3 on the viral genome, promoting heterochromatin formation (19, 20). Maintenance of heterochromatin makes sense for viral latency, in which transcription is restricted, and the DNA is assembled into nucleosomal heterochromatin (46, 47). In contrast, histone association with the viral genome is low during lytic infection (8, 33, 34), and the viral chromatin structure is heterogeneous (48, 49). ATAC-Seq studies have shown that the viral genome is highly accessible during lytic infection (34), with no evidence for nucleosome-mediated protection of DNA (42). ATRX knockout also did not significantly affect genome accessibility in the first 4 h of infection (42). Further suggesting a lack of heterochromatin, PRO-Seq studies from our laboratory have shown that the viral genome is highly transcriptionally active throughout lytic infection (3, 5, 22, 27).

Taken together, these findings suggest that the genome is not incorporated into heterochromatin during early lytic infection. Though H3.3 is deposited on the viral genome during lytic infection (19, 49, 50), it is important to note that H3.3 is enriched at accessible chromatin regions containing active transcriptional regulatory elements such as promoters and enhancers on the cellular genome (51, 52). In addition, incorporating H3.3 is known to promote transcription on the HSV-1 genome (50, 53). Previously, H3.3 loading at active transcription sites was thought to depend on the histone chaperone HIRA, whereas ATRX function was limited to H3.3 loading at heterochromatin (54). However, recent advancements in ChIP using EGS to preserve indirect and/or hyperdynamic interactions (21) have shown that ATRX is associated with H3.3 binding at active transcriptional regulatory elements and G-rich regions, which are putative G4 forming sites (29).

Our finding that ATRX associates with viral loci exhibiting both high and low transcriptional activity mirrors its diverse roles on the cellular genome. A key feature of genomic regions with ATRX binding at active transcription is the presence of G-rich sequences, characteristic of G4-forming motifs (37). The HSV-1 genome contains several experimentally validated G4 sites (38, 55), most notably at the promoters of highly transcriptionally active IE genes (39). Destabilization of these G4s has been shown to promote IE gene expression (56). Interestingly, ATRX depletion alters the expression of cellular genes with G4 motifs in their promoters (29, 32), a phenomenon thought to result from increased G4 formation. (32, 57).

We hypothesized that ATRX may function similarly on the viral genome by facilitating G4 destabilization and that ligand-induced stabilization of G4s would mimic ATRX loss. Consistent with this model, treatment with the G4-stabilizing ligand, BRACO-19 (also known to exhibit antiviral activity by reducing viral DNA polymerase processivity (38)), phenocopied ATRX depletion. Specifically, G4 stabilization led to a comparable reduction in transcriptional initiation of IE genes (Fig. 9A). Interestingly, as with ATRX depletion, transcription of L/ST, which lacks a G4 motif in its promoter, was not affected by BRACO-19 further supporting the hypothesis that ATRX regulates transcription via its interaction with promoter G4s.

G4 stabilization reduces transcriptional initiation and subsequent nascent RNA synthesis of cellular genes due to impaired loading of general transcription factors, including TATA-binding protein (TBP) (58). Since most HSV-1 promoters contain TATA boxes and TBP is associated with IE promoters (34, 59), our model proposes that enhanced G4 formation - presumably occurring in the absence of ATRX - hinders the recruitment of TBP and other factors necessary for efficient transcription.

Although ATRX knockdown increases G4 formation on cellular DNA (57, 60, 61), directly confirming a similar effect on the viral genome is experimentally challenging. Prior studies primarily relied on immunofluorescence to quantify G4 formation, which cannot distinguish between viral and cellular DNA. However, the striking phenotypic similarity between ATRX depletion and G4 stabilization with respect to impaired viral transcriptional initiation strongly suggests a mechanistic link between ATRX and G4s. Nonetheless, this connection remains to be fully validated experimentally.

Incorporation of our data with previous studies supports a model in which incoming HSV-1 genomes are rapidly entrapped and repressed by PML-NBs. Rather than being purely defensive, this repression may benefit the virus by promoting TIEMR to help coordinate the subsequent transcriptional cascade. As viral genomes lack canonical histones early in infection (8, 34, 49, 62), this repression likely results from DNA compaction rather than canonical heterochromatin formation. Our data indicates that DAXX and ATRX have distinct roles in regulating the early transcription of PML-NB-entrapped viral genomes. DAXX may be repressive by promoting H3.3 incorporation, thus enhancing genome compaction (49). This repression is alleviated only when sufficient levels of ICP0 are expressed to displace DAXX. Here, we show that ATRX facilitates IE transcription (and thus ICP0 expression) of this compacted genome by interacting with the G4-containing IE promoters, thereby helping the virus to proceed with de-repression to progress through the temporal cascade.

Data showing reduced mRNA production from genes across three temporal classes support the hypothesis that ATRX depletion delays the progression of the temporal cascade (Fig. S4A). Although overall transcript levels were lower in the absence of ATRX across the time course, the subsequent rate of production of mRNA between 3–6 h post-infection was significantly higher than in control (Fig. S4B, C). Thus, ATRX may contribute to transcriptional repression during later stages of lytic infection as previously reported (19, 20, 42, 63). Interestingly, most evidence for ATRX-mediated repression during lytic infection comes from studies using ICP0-null viruses. These models may not fully capture the early transcriptional interplay between ICP0 (and other IE genes) and ATRX, especially given that ICP0 mutants exhibit dysregulated early transcription (5).

Given ATRX’s well-established, diverse roles in cellular transcription (29), it is reasonable to propose that ATRX plays complex and dynamic roles in regulating viral gene expression. We suggest that ATRX function is influenced by the chromatin state of the viral genome—acting in a pro-transcriptional capacity on IE genes when the incoming genome is largely free of canonical histones, but shifting to a repressive role as the genome becomes more chromatinized. This model also supports a role for ATRX in maintaining transcriptional repression during latency when the viral genome is enriched in repressive histone modifications such as H3K9me3 (20, 50).

Collectively, our findings highlight the complexity of HSV-1’s transcriptional regulation, shaped over millions of years of co-evolution with humans (64). The virus must balance repression—minimizing immune detection—with activation, ensuring that a subset of genomes initiate productive infection. This finely tuned regulation is likely essential in neurons, where the virus must preserve host cell viability to establish and maintain latency.

## Materials & Methods

### Cells

HEp-2 (human epithelial lung cancer) and Vero (African green monkey) cells were maintained in Dulbecco’s modified Eagle’s medium (DMEM) containing 10% new-born calf serum (NBS), 100 units/ml penicillin, 100μg/ml streptomycin (pen/strep) and maintained at 37□JC with 5% CO_2._ HFF (hTERT immortalized human foreskin fibroblasts, BJ-5ta) cells were acquired from ATCC and maintained in 4:1 DMEM:199V medium supplemented with 10% Foetal bovine serum (FBS), pen/step and 0.01mg/ml hygromycin B.

### siRNA Knockdown

Dicer-Substrate siRNAs were ordered from Integrated DNA Technologies (IDT) (65) for PML (hs.Ri.PML.13.1, #452537617), DAXX (hs.Ri.DAXX.13.1, #467654972), ATRX (hs.Ri.ATRX.13.1, #452537611) and non-targeting negative control (DS NC1, #51-01-14-04). 50nM siRNA was reverse transfected using Lipofectamine RNAimax reagent, and cells were seeded on top in a pen/strep-free medium.

### mRNA RT-qPCR

RNA was extracted from whole cell extracts using Monarch Spin RNA Mini Kit (NEB). Reverse transcription was performed using SuperScript III (Invitrogen), primed with Oligo(dT). qPCR analysis of the cDNA was performed using Luna Universal qPCR Master Mix (NEB). Primers are listed in Table S1. Relative expression to the negative control knockdown was calculated using the ΔΔCt method as below:

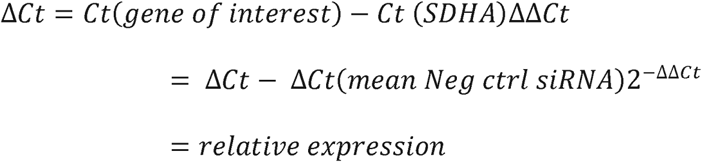

### Viruses, Infection, and Drug Treatment

HSV-1(F) stocks were prepared and titered on Vero cells. Monolayers of HEp-2 cells were infected with HSV-1 at an MOI of 5 in 199V medium supplemented with 1% NBS and pen/strep. Cells were incubated at 4°C for 1 h to allow for virus adsorption. After 1 h, the inoculum was removed and replaced with pre-warmed (37°C) DMEM with 2% serum, and infection was allowed to proceed. This was time 0 hpi. For BRACO-19 treatment, 25μM (or an equal volume of DMSO) was added to the pre-warmed media at 0 hpi.

### Nuclei extraction

Nuclei were isolated from infected using cells as previously described (66, 67). For siRNA knockdowns, 2×10^6^ cells were used per replicate; for all other experiments, 8×10^6^ cells were used. Cells were washed 2x with ice-cold PBS incubated for 10 min on ice with swelling buffer (10mM Tris-HCl [pH 7.5], 10% glycerol, 3mM CaCl_2_, 2mM, MgCl2_2_, 0.5mM DTT, protease inhibitors [Pierce] and 4 U/ml RNase inhibitors [RNaseOUT, ThermoFisher]). Cells were scraped from the plate, pelleted via centrifugation at 600 x g for 10 min (4°C), resuspended in lysis buffer (swelling buffer + 0.5% Igepal), and then incubated on ice for 20 min to release nuclei. Nuclei were pelleted by centrifugation at 1500 x g for 5 min (4°C), washed 2x with lysis buffer with a final wash in storage buffer (50mM Tris-HCl [pH 8.0], 25% glycerol, 5mM MgAcetate, 0.1mM EDTA, 5mM DTT). Nuclei were resuspended in a storage buffer, flash-frozen in LN_2,_ and stored at -80°C.

### Nuclear run-on

For RT-qPCR, frozen nuclei were thawed on ice and a 2-biotin run-on performed by addition of an equal volume of run-on buffer (10mM Tris-HCl [pH 8.0], 5mM MgCl_2_, 1mM DTT, 300 mM KCl, 200μM ATP, 200μM UTP, 200μM biotin-11-GTP, 200μM biotin-11-CTP, 0.4 U/ml RNase inhibitor and 1% Sarkosyl). Run-on was performed under constant shaking for 5 min on a vortex shaker at 37°C.

For PRO-Seq, frozen nuclei were thawed on ice and a 4-biotin run-on was performed as previously detailed (66, 67). An equal volume of run-on buffer (10mM Tris-HCl [pH 8.0], 5mM MgCl_2_, 1mM DTT, 300 mM KCl, 20μM biotin-11-ATP, 20μM biotin-11-GTP, 200μM biotin-11-CTP, 20μM UTP, 0.4 U/ml RNase inhibitor and 1% Sarkosyl) was added to thawed nuclei. Run-on was performed under constant shaking for 3 min on a vortex shaker at 37°C.

All run-on reactions were ended by adding TRIzol LS (ThermoFisher) and RNA extracted following the manufacturer’s protocol (Invitrogen, MAN0000806).

### PRO-RTqPCR

Extracted RNA was subjected to sequential enzyme digestion to ensure the removal of viral DNA, first with XbaI (NEB) for 2h at 37°C and then with TURBO DNase (Invitrogen) overnight at 37°C. RNA was cleaned-up using the Monarch Spin RNA Cleanup Kit (NEB) and biotinylated RNA was purified using streptavidin M280 Dynabeads (Invitrogen), washed 1x in ice-cold high salt wash (50mM Tris-HCl [pH 7.4], 2M NaCl, 0.5% Triton X-100, 0.4 U/ml RNase inhibitor, 1x in ice-cold medium salt wash (10mM Tris-HCl [pH 7.4], 300mM NaCl, 0.1% Triton X-100, 0.4 U/ml RNase inhibitor and 1x in ice-cold low salt wash (5mM Tris-HCl [pH 7.4], 0.1% Triton X-100, 0.4 U/ml RNase inhibitor RNA was eluted from beads via TRIzol extraction and reverse transcribed using SuperScript III (Invitrogen), primed with random hexamers. qPCR analysis of the cDNA was performed using Luna Universal qPCR Master Mix (NEB). Primers are listed in Table S1. Relative run-on activity to the negative control knockdown was calculated using the ΔΔCt method as below:

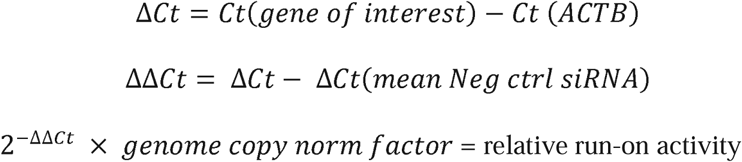

### PRO-Seq Library Preparation

Libraries were prepared following a modified, rapid PRO-Seq protocol (68). RNA was extracted from run-on nuclei via TRIZOL extraction and subjected to base hydrolysis for 10 min on ice with 0.2 NaOH. Unincorporated nucleotides were removed through buffer exchange in a P-30 column (Bio-Rad). As described above, biotinylated RNA was purified using streptavidin M280 Dynabeads (Invitrogen). The 3’-RNA adapter (RA5) was ligated to the 3’ end of the RNA using T4 RNA ssRNA ligase I (NEB).

Biotinylated RNA was again bound to streptavidin M280 beads and 5’ cap removal with 10 U of 5’-pyrophosphohydrolase (RppH) (NEB) and 5’ end repair with T4 PNK (NEB) was performed on beads for 1h at 37°C. Beads were washed 1x high salt, 1x low salt, 1 x DEPC H_2_O between enzyme incubation steps. Ligation of the RA3 5’-RNA adapter was performed on beads using T4 RNA ssRNA ligase I (NEB). RNA was eluted from beads via TRIzol extraction and reverse transcribed with SuperScript III (Invitrogen) using RNA PCR primer 1. The cDNA was PCR amplified with Phusion high-fidelity DNA Polymerase (NEB) using barcoded Illumina PCR index primers. Libraries were purified on an 8% polyacrylamide-TBE gel and underwent paired-end sequencing on an Illumina NovaSeq X Plus (2×150bp) (Novogene, Sacramento, CA, USA).

### PRO-Seq Data analysis

FASTQ files were processed using the PRO-Seq 2.0 pipeline, developed by the Danko lab at Cornell: https://github.com/Danko-Lab/proseq2.0. The genome used to align reads was a concatenated file containing hg38 and HSV-1 F genomes. HSV-1 F genome build had the external repeats deleted to aid sequencing alignment; the modified HSV-1 F genome file is available: https://github.com/Baines-Lab/Public/tree/main/HSV-1. Data were normalized for sequencing depth based on total paired reads and viral genome copy number, detailed in Table S3. HSV-1 normalized bigwig files were visualized using IGV genome browser (69).

### Viral Genome Copy Number Quantification

After nuclear run-on, TRIzol LS RNA extraction, the lower interphase, and organic layers were saved. The DNA was subsequently isolated following the manufacturer’s protocol for TRIzol DNA isolation (Invitrogen, MAN0000806). DNA was cleaned up by 2x phenol:chloroform:isoamyl alcohol purification, followed by ethanol/sodium acetate precipitation and resuspended in DEPC H_2_O. DNA was quantified using Qubit high-sensitivity dsDNA kit. 5ng of DNA was used as input, and qPCR performed using Luna Universal qPCR Master Mix (NEB) with UL51 primers (sequences in Table S1). Genome copy number was determined by standard curve for UL51 as previously described (22, 67). The genome copy norm factor was calculated as below:

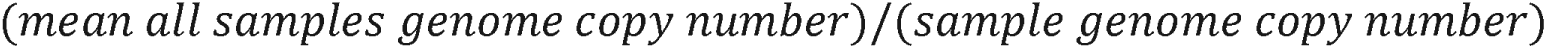

### Immunofluorescence and Confocal Microscopy

siRNA transfected HEp-2 cells seeded onto 13mm glass coverslips were fixed in 4% paraformaldehyde (PFA) at 1.5hpi for 15 minutes, washed in 3XPBS and permeabilized by adding 0.2% Triton X-100 diluted in PBS for 10 minutes. Cells were washed 3XPBS and blocked for 1 h at room temperature in 2% donkey serum (DS) diluted in PBS. Primary antibodies were diluted in PBS + 2% DS and incubated with the cells in a humidity chamber for 1 h. Cells were washed 3X in PBS and incubated with secondary antibodies diluted in PBS + 2% DS in a humidity chamber for 1 h. Cells were washed 3X in PBS, stained with DAPI for 5 min, rinsed 3x in PBS with a final rinse in distilled H_2_O, and mounted onto a microscope slide using ProLong Glass Antifade Mountant (Invitrogen). All incubation steps were performed at room temperature. The details of the antibodies used are in Table S2.

1024×1024 images were acquired on an Olympus Fluoview FV3000 laser scanning confocal microscope using the 60x oil immersion lens. Images were processed in ImageJ Fiji (Version: 2.14.0.154f). A custom CellProfiler (25) pipeline was developed to count nuclear foci in processed images.

### Chromatin Immunoprecipitation and Sequencing (ChIP-Seq)

ATRX-ChIP was performed following the optimized protocol described in (21). In brief, 10^8^ HSV-1 infected (MOI: 5) HEp-2 cells were resuspended in PBS at 1.5hpi and crosslinked with 2mM EGS for 45 min at room temperature, followed by a 1% formaldehyde fixation for 8 min, then quenched in 125mM glycine. Cells were washed 3x in ice-cold PBS and lysed sequentially, first with lysis buffer 1 (100□JmM HEPES, 140□JmM NaCl, 1□JmM EDTA, 10% glycerol, 0.5% NP-40 and 0.25% Triton X-100), then with lysis buffer 2 (200□JmM NaCl, 1□JmM EDTA, 0.5□JmM EGTA and 10□JmM Tris pH 8) and finally with lysis buffer 3 (1□JmM EDTA, 0.5□JmM EGTA, 1□JmM Tris–HCl pH 8, 100□JmM NaCl, 0.1% sodium deoxycholate and 0.5% N-lauroyl sarcosine). All lysis buffers also contained protease inhibitors (Pierce). Samples were sheared with a G27 needle before sonication in a Bioruptor Plus (Diagenode) (45 cycles of 30s on/30s off, 4°C). Sonicated chromatin was pre-cleared with Protein A Dynabeads (Invitrogen) for 1h at 4°C, and 5% (vol) of the sample was saved as input. The remaining sample was split into two and incubated with either Protein A Dynabeads pre-coated with 10 mg anti-ATRX antibody (#ab97508) or 10 μg IgG (Abcam (#ab171870) overnight at 4°C. Beads were washed 5x in wash buffer (50□JmM HEPES, 1□JmM EDTA, 0.7% sodium deoxycholate, 500□JmM LiCl) and bound chromatin eluted with 0.5% SDS and 100 mM sodium bicarbonate. DNA was reverse crosslinked in 0.2M NaCl at 65°C overnight and extracted by phenol:chloroform:isoamyl alcohol purification, followed by ethanol/sodium acetate precipitation and resuspended in DEPC H_2_O.

Sequencing libraries were prepared using NEBNext Ultra II DNA Library Prep Kit for Illumina (#E7645S, NEB), without size selection, following the manufacturer’s protocol. Libraries underwent paired-end sequencing on an Illumina NovaSeq X Plus (2×150bp) (Novogene, Sacramento, CA, USA).

### ChIP-Seq Data Analysis

FASTQ files were processed using a custom pipeline (https://github.com/Baines-Lab/Public/blob/main/ATRX_ChIP_Seq/QC_align.sh) to the same concatenated genome as used in PRO-Seq (hg38 and HSV-1 F). The HSV-1 reads were extracted from the .bam files and normalized using DeepTools (70) bamCoverage with RPGC normalization (-- effectiveGenomeSize 136446) from RPGC normalization of hg38 reads (-- effectiveGenomeSize 2913022398) to generate bigwig files. Peaks were called using MACS3 (30). The bdgdiff command was used to call peaks on the HSV-1 genome (https://github.com/Baines-Lab/Public/blob/main/ATRX_ChIP_Seq/MACS3.sh). HOMER findMotifsGeneome.pl (36) was used to identify enriched motifs in ATRX peaks at IE genes using non-IE peaks as background.

### Statistical Analysis

Statistical analysis was performed in R (version 4.4.2). Details of statistical tests and p-values are detailed throughout.

### Data Availability

The data will be publicly available upon publication on the GEO database under the accession numbers: GSE293672 (access key: qjmbiuyqdvspbop), GSE293673, (access key: olavwiuchdqljat), GSE293674 (access key: cpwvguggjjgzpuj), GSE293675 (access key: chubuawofnyxroz). Reviewers can access the data using keys noted.

## Acknowledgements

We the BRC Genomics Facility (RRID:SCR_021727) at the Cornell Institute of Biotechnology for equipment use. We thank Dr. Claire Birkenheuer for helpful discussions. These studies were supported by National Institutes of Health grants R01 AI 141968 and R21 AI 148926 to J.D.B.

