## Supplementary Figures and Tables for "ATRX promotes transcription initiation of HSV-1 immediate early genes during early lytic infection"

FIG. S1

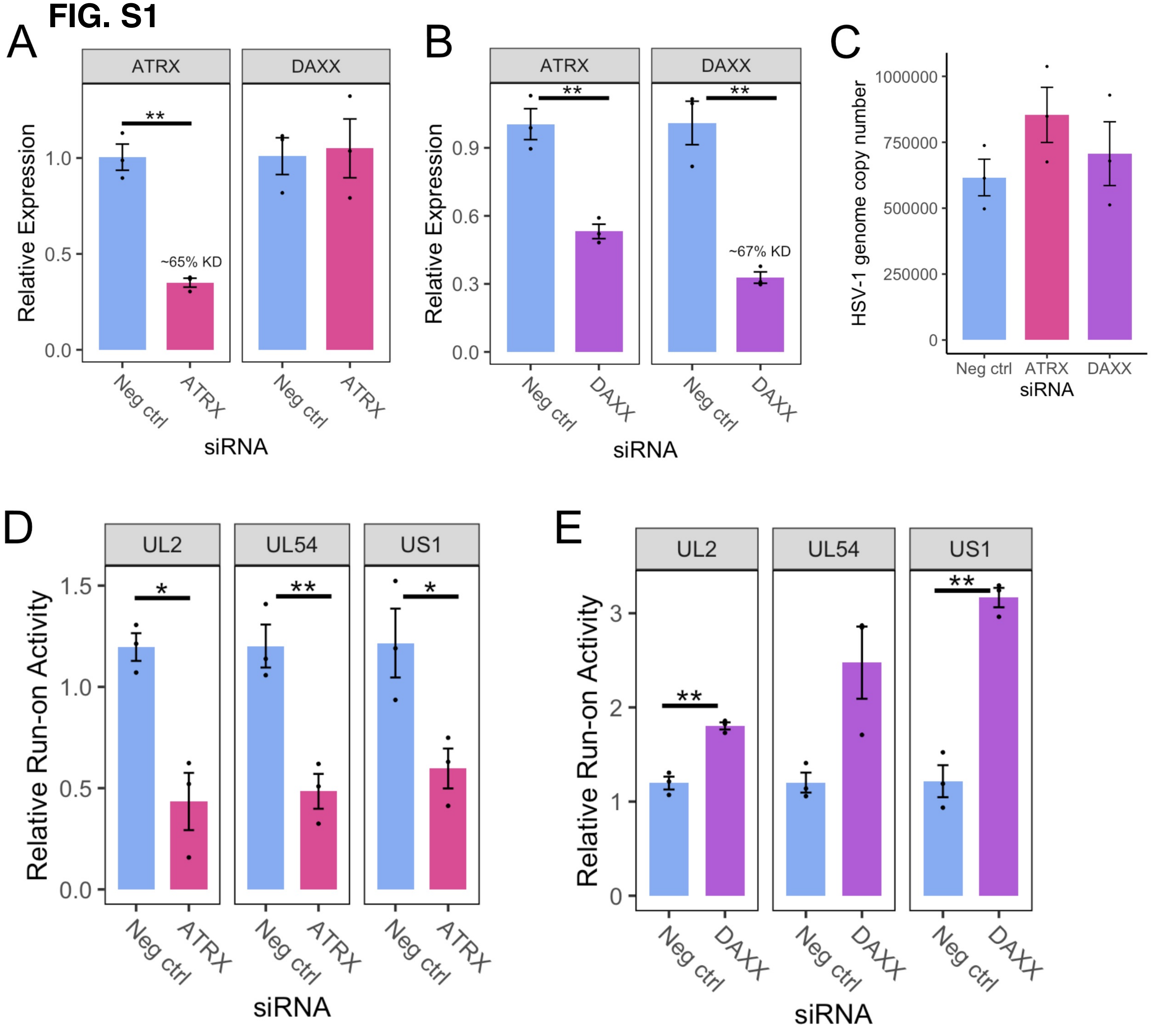


Figure S1: A) RT-qPCR validation of ATRX mRNA KD in HFF cells and DAXX expression. B) RT-qPCR validation of DAXX mRNA KD in HFF cells and ATRX expression. Data are mean +/- standard error. Plotted values are relative to the average negative control (non-targeting), normalized to SDHA. C) HSV-1 genome copy per 5 ng of DNA in siRNA knockdown at 1.5 hpi, determined by UL51 plasmid standard curve qPCR. Data are mean -/+ standard error. Nascent transcriptional activity of HSV-1 genes at 1.5 hpi during ATRX knockdown (D), and DAXX knockdown (E). Data are mean -/+ standard error. Plotted values are relative to the average negative control (non-targeting), normalized to ACTB and viral genome copy number. Statistical significance was determined using Welch's t-test. Asterisks indicate statistical significance (*=p<0.05, **=p<0.01).

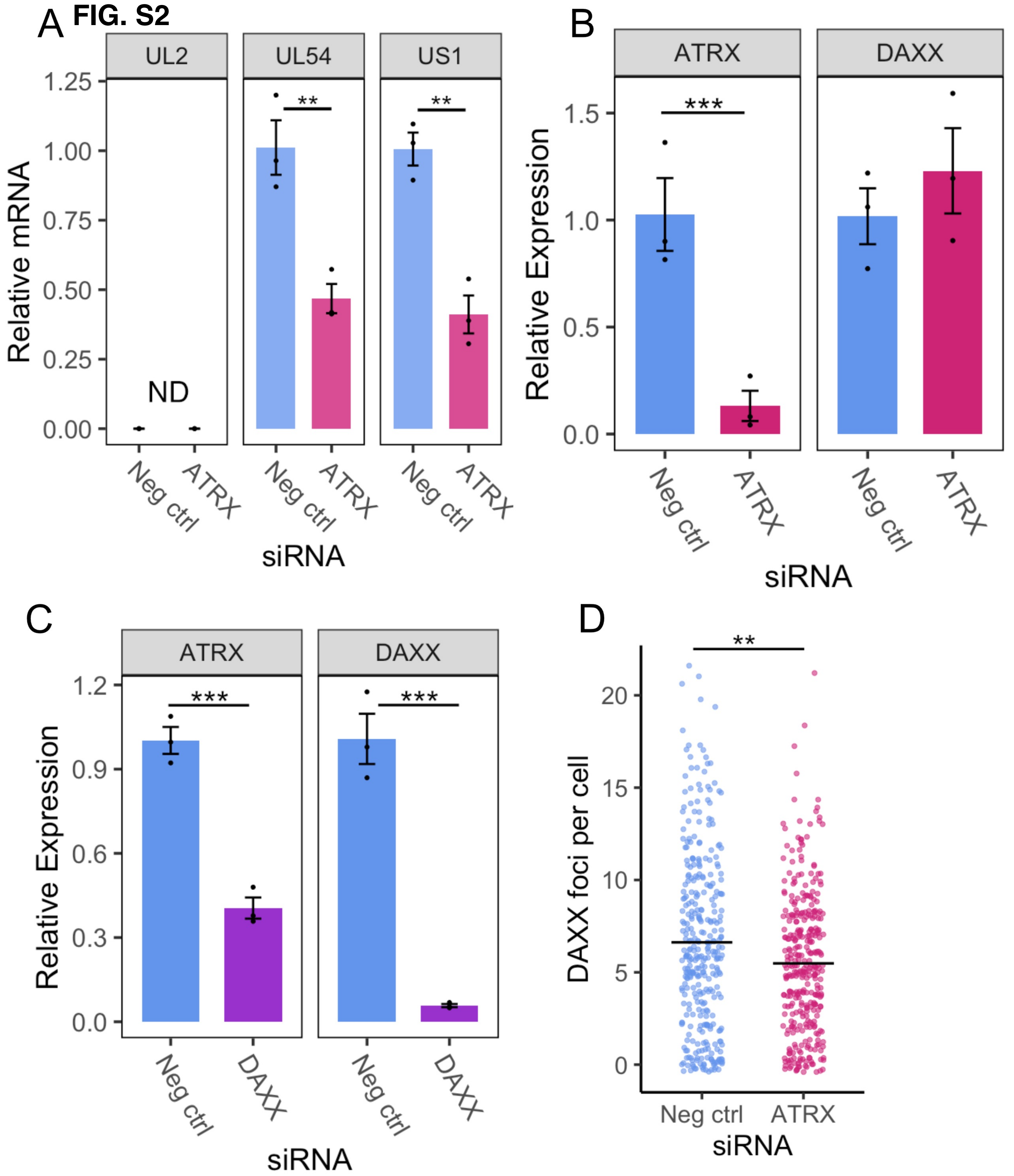


Figure S2: A) RT-qPCR of HSV-1 mRNA during ATRX KD at 1.5 hpi in HEp-2 cells. B) RT-qPCR of ATRX and DAXX mRNA during ATRX KD at 1.5 hpi in HEp-2 cells. ND not detected. C) RT-qPCR of ATRX and DAXX mRNA during DAXX KD at 1.5 hpi in HEp-2 cells. All RT-qPCR Data are mean +/- standard error. Plotted values are relative to the average negative control (non-targeting), normalized to cellular gene SDHA. D) Quantification of DAXX and foci per cell (nucleus) during ATRX KD. Each dot represents an individual cell (**=p<0.01,***=p<0.001).

FIG. S3

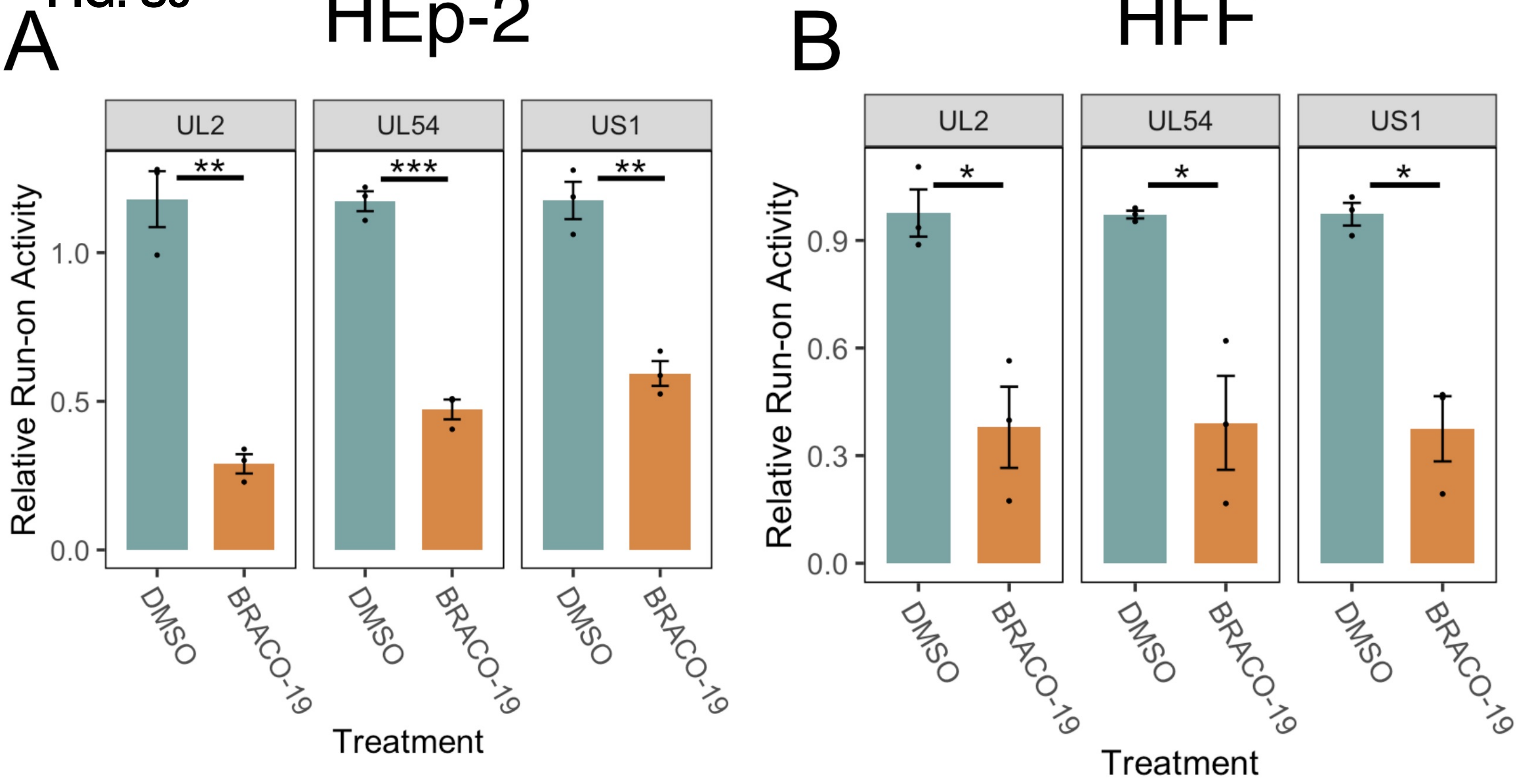


Figure S3: Nascent transcriptional activity of HSV-1 genes at 1.5 hpi during BRACO-19 treatment in HEp-2 (A) and HFF (B) cells determined by PRO-qPCR. Data are mean -/+ standard error. Plotted values are relative to the average negative control (non-targeting), normalized to ACTB and viral genome copy number. Statistical significance was determined using Welch's t-test. Asterisks indicate statistical significance (*=p<0.05, **=p<0.01, ***=p<0.001).

**FIG. S4**

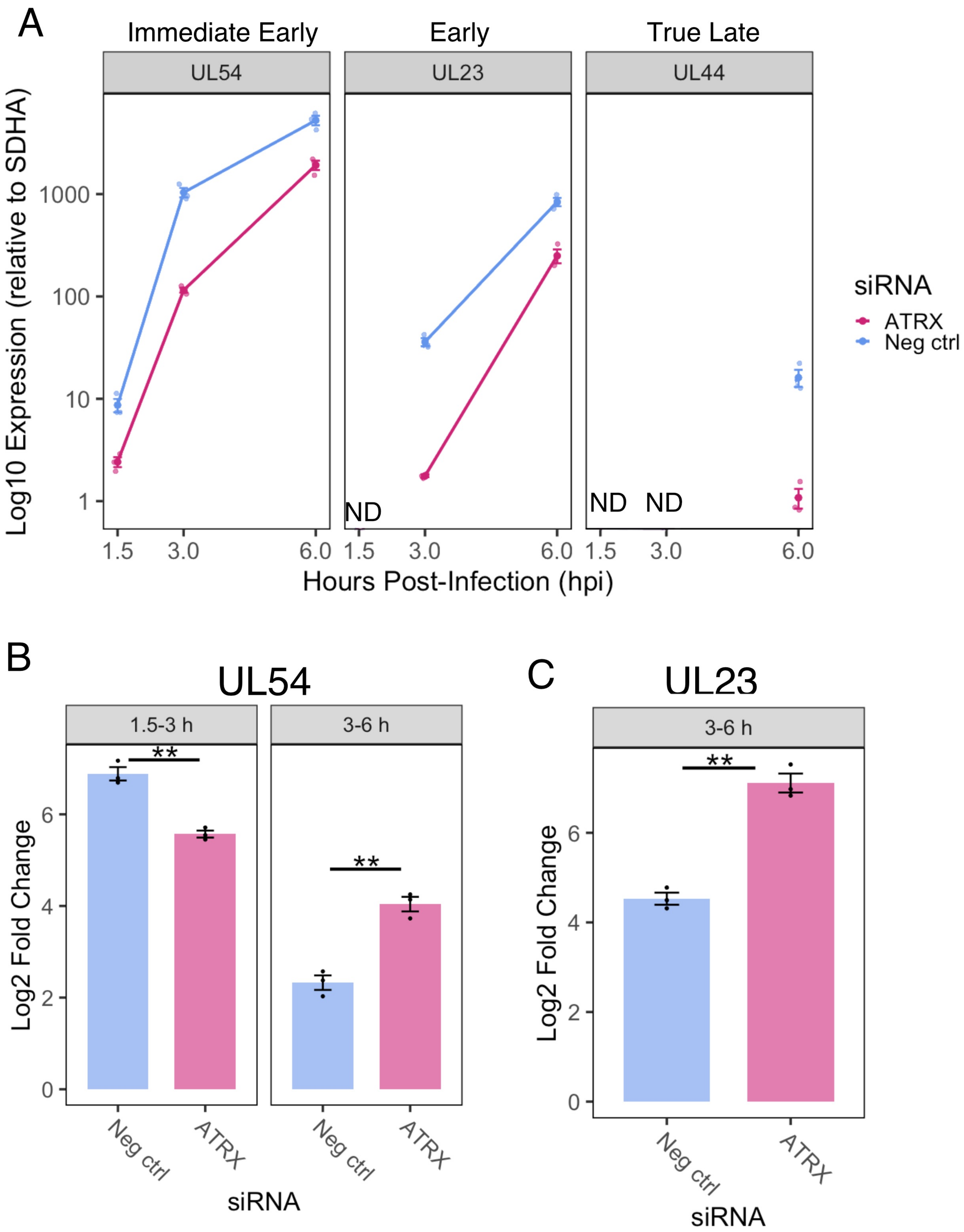


Figure S4: A) Relative HS V-1 mRNA (normalized to SDHA) of three temporal classes at 1.5, 3 and 6 hpi. ND = not detected. Fold change in expression from 1.5-3 and 3-6 hpi for UL54 (B) and from 3-6 hpi for UL23 (C). Data are mean +/ - standard error. Statistical significance was determined using Welch's t-test. Asterisks indicate statistical significance （*=p<0.05， **=p<0.01）

**Table S1: Primers**

| Gene | Forward (5'-3') | Reverse (5'-3') | Target |
|---|---|---|---|
| **PML** | CCGTCATAGGAAGTGAGG CTTC | GTTCGCATCTGAGTCTCCG | mRNA |
| **ATRX** | ACGGCGTAGTGGTTGTCTC | GCAGCATGTAGCCCTCT | mRNA |
| **DAXX** | GAAGCTCTGGATTCTGTG | CATCACTCTCTCATCGTCTCG | mRNA |
| **SDHA** | GGACAACTGGAGGTGGCATT | CCGTCATGTAGTGGATGGCA | mRNA |
| **UL23** | ACCGCTAACAGCGTCAACA | CAAAGAGGTGCGGGAGTT | mRNA |
| **UL44** | GTGACGTTTGCCTGGTTCCTGG | GCACGACTCCTGGCCGTACG | mRNA |
| **UL2** | CTGTGAAGGCTGGGTGTG | GTAGT CAAATCTGTCGC | Nascent RNA/mRNA |
| **UL54** | GCATCCTTCGTGTTTGTCATTCT G | GCATCTTCTCTCCGACCCCG | Nascent RNA/mRNA |
| **US1** | TTTGGGGAGTTTGACTGGAC | CAGACACTTGCGGTCTTCTG | Nascent RNA/mRNA |
| **ACTB** | GCTCATGTAGAAGGTGTG | GCATGGTCAGAAGGATCT | Nascent RNA |
| **UL51** | GCCAGTCGTTCTAGGTTCAC | GTTAACGCGCTACTTCCCG | HSV-1 DNA |

**Table S2: Antibodies**

| **Antibody/Stain** | **Source** | **Use** |
| --- | --- | --- |
| **ATRX (Rabbit Polyclonal)** | Bethyl (#A301-045A) | IF (1:100) |
| **DAXX (Rabbit Polyclonal)** | MilliporeSigma (#07-471) | IF (1:100) |
| **PML (Mouse Monoclonal)** | Abcam (#ab9605) | IF (1:500) |
| **Donkey Anti-rabbit IgG Alexafluor 568** | Invitrogen (#A10042) | IF (1:500) |
| **Donkey Anti-mouse IgG Alexafluor 647** | Invitrogen (#31571) | IF (1:500) |
| **DAPI** | Invitrogen (#D1306) | IF (1μg/ml) |
| **ATRX (Rabbit Polyclonal)** | Abcam (#ab97508) | ChIP (10μg per IP) |
| **IgG (Rabbit Polyclonal)** | Abcam (#ab171870) | ChIP (10μg per IP) |

**Table S3: PRO-Seq Normalization Details**

**ATRX KD PRO-Seq**

| *Sample* | *Total Paired Reads* | *Library correction factor* | *HSV1 genome copies (qPCR)* | *Genome copy correction factor* | *Combined correction factor* |
|---|---|---|---|---|---|
| *Neg_KD_1* | 12664906 | 1.12 | 382102.09 | 0.98 | 1.09 |
| *Neg_KD_2* | 13876038 | 1.02 | 352869.89 | 1.06 | 1.08 |
| *Neg_KD_3* | 11975852 | 1.18 | 280353.42 | 1.33 | 1.57 |
| *ATRX_KD_1* | 12083661 | 1.17 | 371827.00 | 1.00 | 1.18 |
| *ATRX_KD_2* | 15752967 | 0.90 | 448850.98 | 0.83 | 0.75 |
| *ATRX_KD_3* | 18534405 | 0.76 | 403396.05 | 0.93 | 0.71 |
| **MEAN** | 14147972 | **MEAN:** | 373233.24 | | |

**HEp-2 1.5 hpi PRO-Seq (for ATRX ChIP-Seq comparison)**

| *Sample* | *Total Paired Reads* | *Library correction factor* | *HSV1 genome copies (qPCR)* | *Genome copy correction factor* | *Combined correction factor* |
|---|---|---|---|---|---|
| *Hep2_1* | 64737291 | 1.07 | 26449.9021 | 1.02 | 1.09 |
| *Hep2_2* | 74188010 | 0.94 | 27429.3679 | 0.98 | 0.92 |
| **MEAN** | 69462650.5 | **MEAN** | 26939.635 | | |

**BRACO-19 PRO-Seq**

| *Sample* | *Total Paired Reads* | *Library correction factor* | *HSV1 genome copies (qPCR)* | *Genome copy correction factor* | *Combined correction factor* |
|---|---|---|---|---|---|
| *DMSO_1* | 63318485 | 1.06 | 488329.607 | 0.99 | 1.04 |
| *DMSO_2* | 68306699 | 0.98 | 431937.951 | 1.11 | 1.09 |
| *DMSO_3* | 59719519 | 1.12 | 393094.074 | 1.22 | 1.37 |
| *BRACO19_1* | 66106799 | 1.01 | 523448.398 | 0.92 | 0.93 |
| *BRACO19_2* | 78368896 | 0.86 | 512761.915 | 0.94 | 0.80 |
| *BRACO19_3* | 66530182 | 1.01 | 537273.125 | 0.90 | 0.90 |
| **MEAN** | 67058430 | **MEAN:** | 481140.845 | | |
